# Theta-band brain synchronization supports the immediate and post-sleep dynamics of memory recall in children

**DOI:** 10.64898/2026.04.30.721844

**Authors:** Soléane Gander, Coralie Rouge, Anna Peiffer, Philippe Peigneux, Peter Simor, Mathieu Bourguignon, Vincent Wens, Xavier De Tiège, Gaétane Deliens, Charline Urbain

## Abstract

Memory engrams emerge from dynamic, coordinated interactions among synchronized functional brain networks. Yet, how these networks are selectively reactivated upon cued recall and gradually (re)organized over time, especially through sleep, remains poorly understood. Using magnetoencephalography (MEG), sleep electroencephalography (EEG) and behavioral measures, we investigated the spatiotemporal neural dynamics of cued recall memory during immediate and post-sleep (i.e., 90-minute post-learning nap) recall sessions in school-aged children. Results showed that immediate recall engaged a temporally ordered sequence of theta-band phase synchronization across two transiently synchronized networks: an early (150–350 ms) network involving the ventral visual pathway and left medial temporal lobe (MTL), followed by a later (550–750 ms) network encompassing the bilateral MTL and widespread neocortical associative regions. Post-sleep recall was associated, relative to wakefulness, with strengthened theta-band phase synchronization between the left MTL and widespread bilateral neocortical regions in a similar late window (450–650 ms). Post-sleep theta-band synchronization and memory gains in performance positively correlated with slow oscillation–spindle coupling during the post-learning nap. Altogether, these findings highlight oscillatory and spatiotemporal dynamics of memory recall networks and suggest that sleep, possibly driven by slow-oscillation–spindle coupling mechanisms, supports the efficient reinstatement of memory traces in the developing brain.

## 1. Introduction

Alongside genetically determined processes, learning experiences play a central role in shaping neurodevelopment through neural plasticity mechanisms ^1^. In particular, childhood is associated with profound transformations of memory systems ^2^, including the formation and reorganization of engrams, i.e., the physical traces of memory in the brain ^3^. Yet, the neurophysiological mechanisms supporting the dynamic reactivation and reorganization of post-learning memory engrams, that are fundamental processes to cognitive development, remain unknown in children and poorly understood in adults.

Over the past decades, research in adults has primarily relied on functional magnetic resonance imaging (fMRI) to characterize detailed maps of the brain regions and large-scale networks involved in memory processes. Beyond the medial temporal lobe (MTL), long recognized as a critical hub for associative memory ^4,5^, fMRI studies have consistently shown that memory encoding and recall processes engage distributed and MTL-dependent bilateral neocortical networks ^6,7^. However, research focusing on the temporal organization and dynamic interactions of memory networks remains scarce. To date, related insights have primarily emerged from intracranial electroencephalography (iEEG) studies in epilepsy patients. These studies have contributed to a chronometric modelling of cued recall^8^, proposing that memory recall unfolds as a rapid and temporally ordered sequence of reactivation processes across sensory, MTL, and neocortical regions. According to this framework, sensory information is rapidly conveyed to MTL structures after a stimulus presentation, enabling cue-driven recognition within ∼500 ms. This is followed by pattern completion, reflected in the reactivation of hippocampal encoding-related neural assemblies ^9–11^, which subsequently drives the reinstatement of distributed neocortical representations between 500 and 1500 ms after cue onset ^12,13^, itself likely supported by theta-band (4–8 Hz) synchronization that plays a critical role in MTL–neocortical communication ^14–19^.

Critically, although the dynamics of memory recall and associated cortical reinstatement have been investigated in a limited number of non-invasive EEG and magnetoencephalography (MEG) studies in healthy adults ^20–22^, these processes remain unexplored in children. Moreover, to our knowledge, no study has yet characterized the spatiotemporal neural dynamics of memory recall in light of inter-areal theta-band synchronization at the whole brain level, nor its (re)organization over time. This gap is particularly striking given that memory networks not only build up and evolve at subsecond timescales but continue developing over longer time windows, particularly through post-learning offline plasticity periods such as sleep.

In particular, non-rapid eye movement (NREM) sleep has been proposed to promote memory consolidation by progressively strengthening and reorganizing memory traces from the MTL to distributed neocortical regions (see reviews ^23–27^). Converging evidence further suggests that this reorganization depends on the precise temporal coupling between slow oscillations (<1 Hz), thalamocortical sleep spindles (∼11–16 Hz), and hippocampal ripples (∼80 Hz), which together support hippocampal memory reactivation and redistribution within long-term neocortical storage sites ^23,25–33^. Consistent with this framework, precisely timed slow-oscillation–spindle coupling has been shown to promote memory consolidation performance across the lifespan ^34–37^.

Notwithstanding recent advances, only a small number of studies have examined how sleep shapes the spatiotemporal neural dynamics of memory networks. In adults, fMRI studies have reported decreased hippocampal and increased neocortical engagement at recall following post-learning sleep ^38–41^. In children, evidence remains extremely limited, with a single MEG study reporting enhanced recall-related evoked magnetic responses in prefrontal and bilateral inferior parietal regions following sleep ^42^. Yet, this question is particularly salient at school age, since this developmental period is not only marked by heightened neuroplasticity ^1^, but also by an increased slow-oscillation activity during NREM sleep ^43^, two factors likely to amplify sleep-dependent memory consolidation ^44–47^. The extent to which post-learning sleep modulates theta-band synchronization during memory recall, and the contribution of sleep-specific oscillatory mechanisms such as slow-oscillation–spindle coupling, remain largely unexplored.

The present study aims at filing these gaps by characterizing the spatiotemporal dynamics of theta-band synchronization underlying memory recall, and the specific contribution of sleep-dependent processes in the (re)organization of memory-related networks at school age (7–12 years old). We hypothesized that memory recall would rely on transient theta-band synchronized networks, ultimately leading to MTL–neocortical reinstatement processes. Furthermore, we expected that post-learning sleep would enhance memory reinstatement by increasing MTL–neocortical theta-band synchronization during delayed recall, relative to wakefulness. Finally, we predicted that sleep-specific oscillatory activities, particularly slow-oscillation–spindle coupling, would correlate with better recall performance and enhanced post-sleep theta-band synchronization.

## 2. Results

### 2.1. Memory recall performance

Thirty-one children (mean age ± SD: 9.90 ± 1.27 years; range, 7–12 years), randomly assigned to a Sleep (N=15) or Wake (N=16) group completed an associative memory task involving 50 non-object–function associations (see “Methods”). During the learning session, children successfully learned 75 ± 7% of the associations (mean ± SD; range: 60–92%). Sleep and Wake groups did not differ in the number of trials required to reach the learning criterion (≥60% correct) or in overall learning performance (independent-samples t-tests, all *ps* > 0.05), confirming comparable learning levels across groups. Memory recall performance was computed during two cued recall sessions: an immediate recall occurring after learning and a delayed recall following a 90-min nap (Sleep group) or an equivalent period of wakefulness (Wake group). Children were presented with a non-object cue and were asked to verbally recall its associated function. A mixed ANOVA on memory recall performance with session (immediate vs. delayed recall) as a within-subject factor and group (Sleep vs. Wake) as a between-subject factor revealed a main session effect (F(1,29) = 12.08, *p* = 0.002), reflecting better performance during delayed relative to immediate recall, and a main group effect (F(1,29) = 10.1, *p* = 0.004), with higher recall in the Sleep group compared to the Wake group. No significant session × group interaction effect was observed (F(1,29) = 0.02; *p* = 0.877; see supplementary Figure S1 in Supplementary Information, SI1), indicating that the improvement in performance from immediate to the delayed recall was comparable across groups. Consistent with this result, the memory performance gain from immediate to delayed recall did not differ between groups (t(29) = -0.398; *p* = 0.694). Importantly, memory recall effects could not be attributed to differences in vigilance, as no significant effects of group, session or interaction were observed on reciprocal reaction times (RRTs) ^48^ in the psychomotor vigilance task (PVT; all *ps* > 0.05) held before each recall session, and no significant correlations were found between RRTs and memory recall performance (all *ps* > 0.05).

### 2.2. Theta-band synchronization dynamics during immediate memory recall

To characterize the theta-band synchronization dynamics associated with memory recall, MEG data were collected as children performed an immediate cued recall of the 50 non-object–function associations (data combined across Sleep and Wake groups). Immediate recall of the newly learned associations evoked theta-band phase synchronization that peaked in two distinct time windows, in line with Staresina & Wimber (2019) model ^8^, i.e., 150–350 ms and 550–750 ms post-cue onset windows (Figure 1A). To determine the synchronized networks engaged during these two time windows, paired t-tests compared theta-band phase synchronization during each time window with the pre-stimulus baseline (-200 to 0 ms), using the Network Based Statistic software (NBS) ^49,50^. In the early 150–350 ms post-cue window, transient phase synchronization involved a visual pathway–hippocampal network encompassing bilateral occipital and temporal regions within extra-striate visual ventral pathway and the left hippocampus (*p*^corr^ < 0.001; Figure 1B). Hubs within this early network included the right superior occipital gyrus, bilateral calcarine sulcus and the right inferior temporal gyrus. In the later 550–750 ms post-cue window, transient phase synchronization emerged in a widespread MTL–neocortical network comprising bilateral MTL regions and widespread neocortical associative areas, including the right occipital-parietal cortex, the left lingual gyrus, the right inferior temporal gyrus, the right temporal pole, and the right inferior frontal gyrus (IFG) (*p*^corr^ < 0.001, Figure 1C). The full list of nodes, related node strength and MNI coordinates are provided in Supplementary Information, SI2 & SI3.

**Figure 1.**
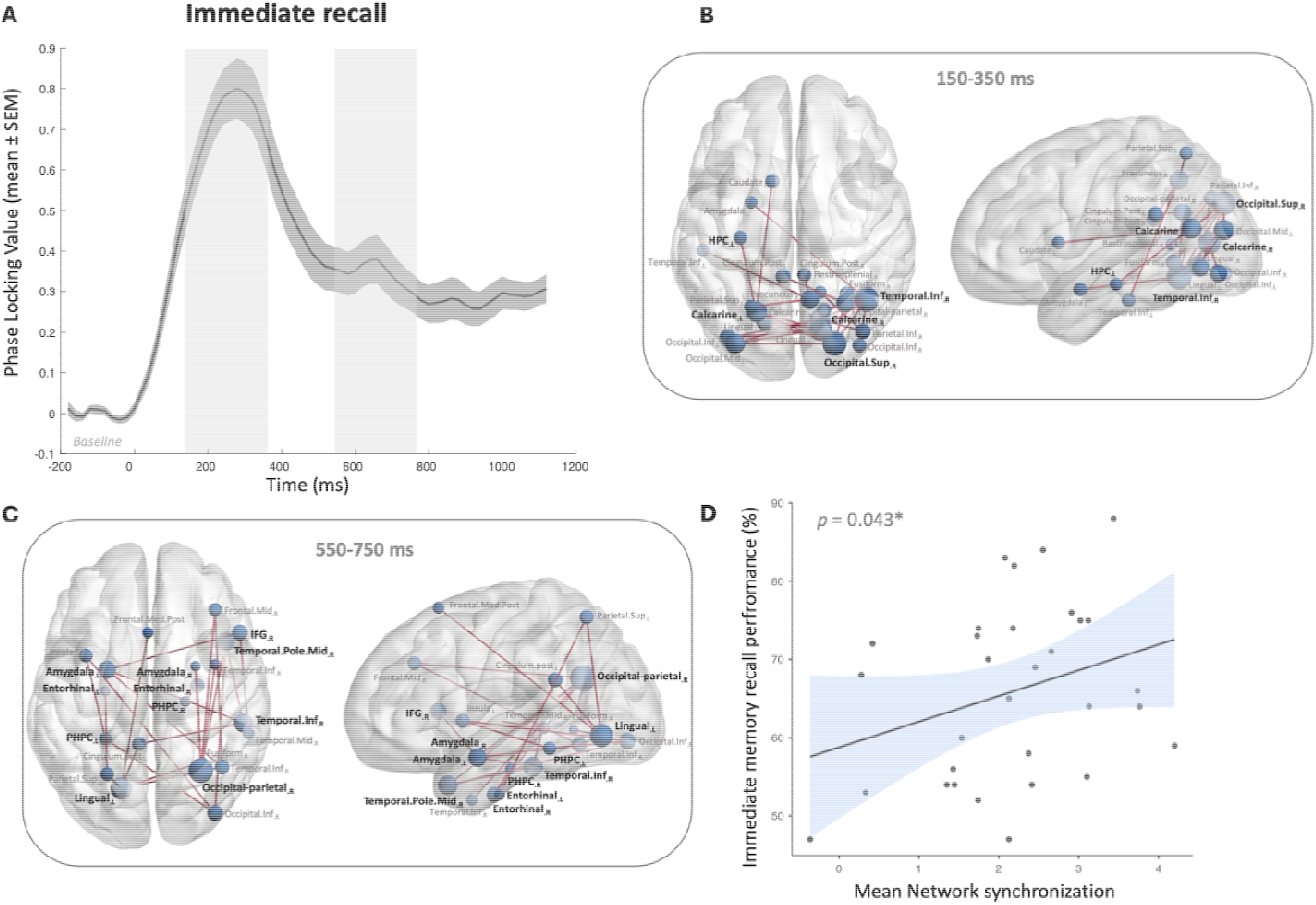
Theta-band synchronization dynamics during immediate memory recall. **(A)** Theta-band phase synchronization computed as cross-trial phase locking value (PLV; mean ± SEM) across all participants (Sleep and Wake groups combined) during the immediate recall session, peaking in two distinct time windows (i.e., 150–350 ms and 550–750 ms post-cue windows). **(B)** Significant synchronized network in the early 150–350 ms post-cue window, compared to the baseline (-200–0 ms). Node size represents node strength (i.e., sum of all weighted connections to a given node). **(C)** Significant synchronized network in the later 550– 750 ms post-cue window, compared to the baseline (-200–0 ms). **(D)** Significant correlation between the mean network synchronization strength in the second network (550–750 ms post-cue window) and immediate memory recall performance. The regression line indicates the best linear fit, and the shaded area around the line represents the standard error of the estimate.

We next sought to characterize the association between these transiently synchronized networks and immediate memory recall performance. No significant correlation was observed with the mean network synchronization strength in the first network (one-tailed *p* = 0.094), but the mean network synchronization strength in the second network positively correlated with immediate memory recall performance (r = 0.313, one-tailed *p* = 0.043; Figure 1D), suggesting that increased theta-band synchronization from 550 to 750 ms within this MTL–neocortical network is linked to better memory recall performance in children.

### 2.3. Impact of sleep on theta-band synchronization dynamics during delayed memory recall

To investigate how sleep affects theta-band synchronization during memory recall, MEG data from the Sleep and Wake groups were compared during immediate and delayed recall after a 90-min nap or an equivalent period of wakefulness. Delayed recall evoked transient theta-band phase synchronization that was increased in the Sleep group as compared to the Wake group in a 450–650 ms post-cue window, an effect that was not observed during immediate recall (Figure 2A). Between-group differences in the synchronized networks supporting recall were assessed using NBS t-tests on theta-band phase synchronization in the 450–650 ms post-cue window. The sleep-dependent increase in theta-band phase synchronization during delayed recall involved a predominantly left-lateralized network encompassing left MTL structures and distributed neocortical regions (*p*^corr^ = 0.023; Figure 2B). Network Hubs included the right inferior occipital gyrus, left postcentral gyrus, and left superior temporal gyrus as well as in the right IFG, the bilateral amygdalae, the left parahippocampal, entorhinal, and perirhinal cortices. The full list of nodes, related node strength and MNI coordinates are provided in Supplementary Information, SI4. Importantly, no equivalent between-group differences were observed prior to sleep during the immediate recall (*ps*^corr^ > 0.05).

**Figure 2.**
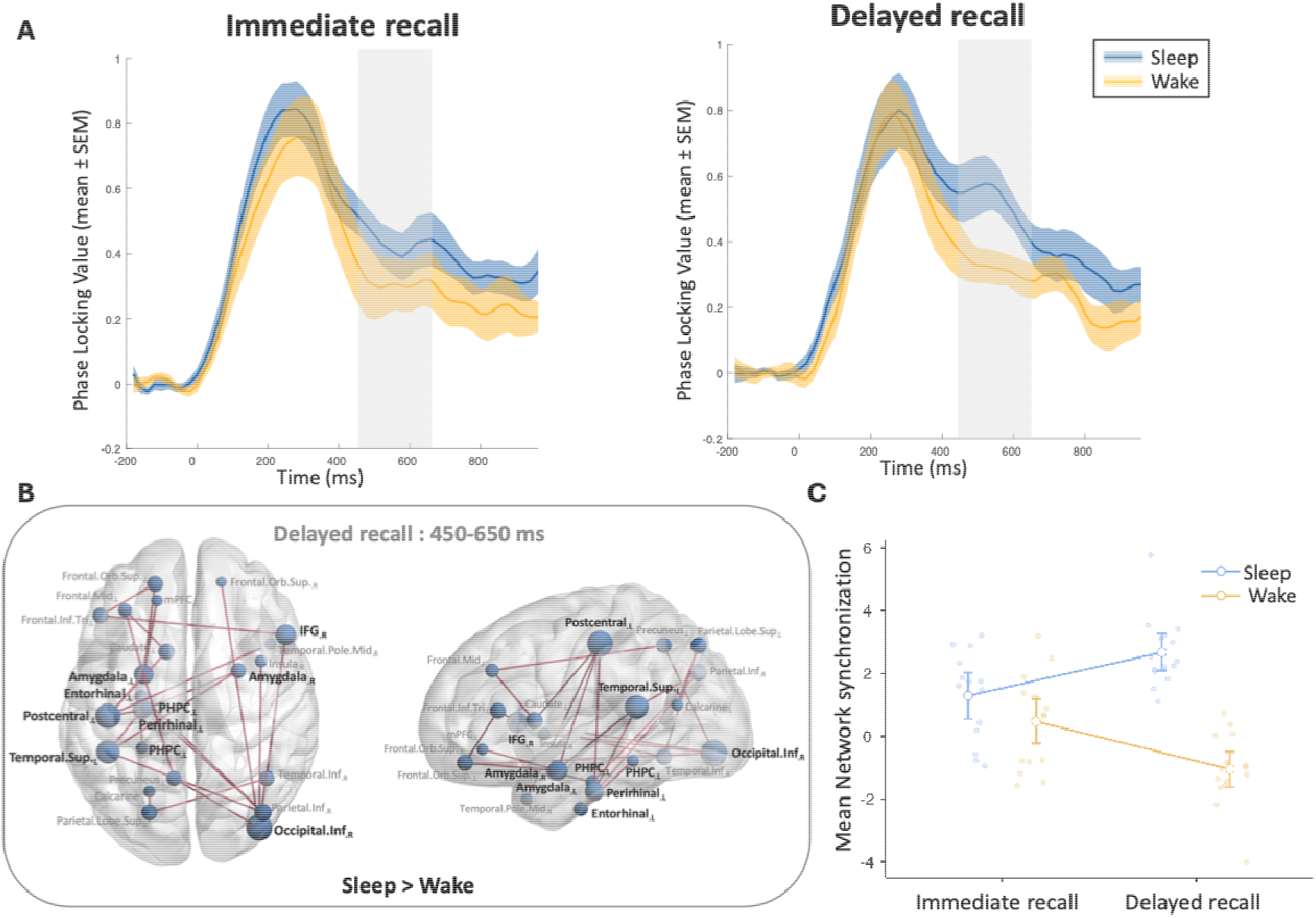
Impact of sleep on theta-band synchronization dynamics during memory recall. **(A)** Theta-band phase synchronization computed as cross-trial PLV (mean ± SEM) increased during delayed recall in the Sleep group (blue) relative to the Wake group (yellow) in the 450–650 ms post-cue window (right panel). During immediate recall (left panel), there was no difference in theta-band phase synchronization (cross-trial PLV; mean ± SEM) between the Sleep group and the Wake group in the 450–650 ms post-cue window. **(B)** Between-group differences in theta-band phase synchronization during delayed recall (Sleep > Wake) in the 450–650 ms post-cue window, revealing a significant synchronized network. Node size represents node strength differences between groups (Sleep > Wake). **(C)** Mean network synchronization strength within this significant sleep-dependent network (450–650 ms) for each group and session, analyzed using a mixed ANOVA. Error bars represent 95% confidence intervals.

We next investigated how sleep impacted the synchronization strength of this sleep-dependent network by comparing mean network synchronization strength across sessions (immediate vs. delayed recall) and groups (Sleep vs. Wake) using a complementary mixed ANOVA (Figure 2C). Results revealed a significant main group effect (F(1,29) = 41.80, *p* < 0.001), no session effect (F(1,29) = 0.06, *p* = 0.809) and a significant group × session interaction effect (F(1,29) = 25.52, *p* < 0.001), explained by an increase in mean network synchronization strength from immediate to delayed recall in the Sleep group (*p* = 0.014), at variance with a decrease between the immediate and the delayed recall in the Wake group (*p* = 0.004).

### 2.4. Associations between slow-oscillation–spindle coupling, post-sleep theta-band synchronization and memory performance

To further investigate how sleep-specific oscillatory activity relates to successful delayed memory recall, we performed correlation analyses between individual measures of slow-oscillation–spindle phase–amplitude coupling (PAC) during the post-learning nap and both memory performance gains and post-sleep network synchronization strength. We first characterized the post-sleep delayed recall network in the Sleep group by comparing theta-band phase synchronization in the 450–650 ms post-cue window with the pre-stimulus baseline (Figure 3A). This analysis revealed a transient increase in theta-band synchronization in the 450–650 ms post-cue window within a predominantly left-lateralized occipito-parieto-temporal network (*ps*^corr^ = 0.013), encompassing the left fusiform gyrus, the left postcentral gyrus, and primarily left MTL areas (Figure 3B). The full list of nodes, related node strength and MNI coordinates are provided in Supplementary Information, SI5. Individual measures of slow-oscillation–spindle PAC during the post-learning nap were positively correlated with the mean network synchronization strength of this post-sleep delayed recall network (r = 0.48, one-tailed *p* = 0.035; Figure 3C), as well as with memory performance gains from immediate to delayed recall (r = 0.57, one-tailed *p* = 0.014: Figure 3D). No comparable correlations were observed between slow-oscillation–spindle PAC and theta-band phase synchronization during immediate recall, nor with immediate recall performance. Of note, this post-sleep delayed recall network did not overlap with the immediate recall network in the Sleep group in the similar 450–650 ms post-cue window (see Supplementary Information, SI6).

**Figure 3.**
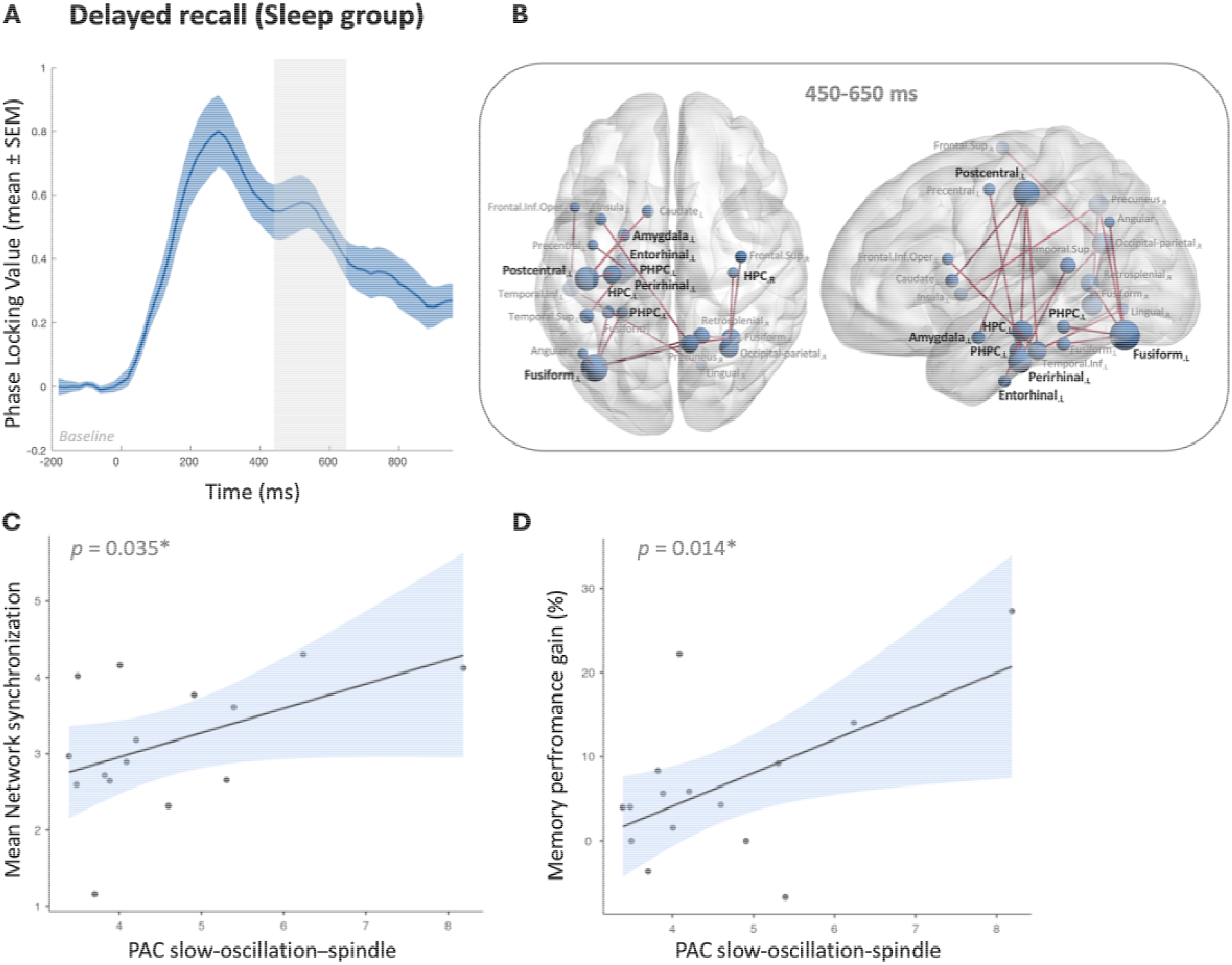
Associations between slow-oscillation–spindle PAC, post-sleep theta-band synchronization and memory performance gain. **(A)** Theta-band phase synchronization computed as cross-trial PLV (mean ± SEM) in the sleep group during the delayed recall session. **(B)** Significant synchronized network in the 450–650 ms post-cue window, compared to the baseline (-200–0 ms). Node size represents node strengths, and hub regions are labelled in bold on the brain plot. **(C)** Significant correlation between the mean network synchronization strength and individual measures of slow-oscillation-spindle PAC. **(D)** Significant correlation between the memory performance gain from immediate recall to delayed recall and individual measures of slow-oscillation-spindle PAC. The regression line indicates the best linear fit, and the shaded area around the line represents the standard error of the estimate.

Importantly, neither total sleep duration nor the duration or proportion of individual sleep stages (see Supplementary Information, SI7) significantly correlated with mean network synchronization strength or memory performance gains (all *ps* > 0.5). Together, these findings suggest a specific role of slow-oscillation–spindle PAC in strengthening post-sleep theta-band phase synchronization and promoting gains in delayed memory recall performance.

## 3. Discussion

By combining whole-brain, noninvasive MEG with sleep EEG, we characterized here the spatiotemporal dynamics of theta-band phase synchronization underlying successful memory recall, and revealed how a post-learning nap specifically modulates related networks in school-aged children, a population marked by heightened neuroplasticity ^1^.

During the immediate recall session, two temporally dissociable theta-band synchronization networks were identified. The spatiotemporal pattern of the early visual pathway-hippocampal network (150–350 ms post-cue) closely mirrors the early (< 500 ms) engagement of sensory and hippocampal areas reported during cued recall, supporting rapid processing of the sensory cue (i.e., the visual non-object) and its familiarity assessment ^8,10,11^. The second MTL–neocortical network (550–750 ms) aligns with neural processes emerging after 500 ms that support the cortical reinstatement of the full memory trace ^12,13,21,22^. Indeed, this later network was not only recruited at a relevant time window (550–750 ms) for memory reinstatement processes but also involved extensive MTL–neocortical functional connections, including bilateral entorhinal and parahippocampal cortices, known to support the hippocampal–neocortical communication critical for associative memory retrieval ^51–53^. Accordingly, this later network encompassed functional connections between key nodes of the ventral visual pathway, likely supporting the retrieval of stored visual non-object representations ^54–57^, and the right IFG, right middle temporal gyrus, and right temporal pole, known to trigger the controlled access to associated semantic representations (i.e., the non-object’s magical function) ^58,59^. Bilateral amygdala involvement was also observed, potentially reflecting arousal-related mechanisms that facilitate the retrieval and/or the anticipation of successful memory recall ^60,61^. These findings suggest that this later distributed network (> 500 ms post-cue) supports the reinstatement of the full memory trace, enabling recall of the learned associations between visual non-objects and their corresponding semantic functions. Last but not least, the network synchronization strength of this later network positively correlated with immediate recall performance, highlighting the relevance of large-scale MTL–neocortical synchronization for the successful reinstatement of the full memory trace. Together, these post-learning MEG findings closely align with a current model of cued recall chronometry mostly derived from iEEG epileptic adults’ studies, which propose sequential dynamics from early sensory processing and rapid MTL engagement within the first ∼500 ms, to later neocortical reinstatement supporting the recall of associative memory traces ^8^. Extending this framework, our results demonstrate that associated theta-band phase synchronization dynamics can be captured noninvasively at the whole-brain level in a pediatric, typically developing population, and suggest a link between reinstatement-related network and recall success.

We next examined how a post-learning nap modulates memory recall-related brain networks, as identified by theta-band phase synchronization. MEG results showed that, as compared with wakefulness, a 90-minute nap enhanced theta-band synchronization across a widespread, predominantly left-lateralized network encompassing left MTL structures and distributed neocortical regions. Notably, this sleep-dependent effect emerged within the predefined critical time window for memory reinstatement (450–650 ms post-cue onset) ^8^, and no comparable between-group difference was observed during the immediate recall session, before sleep. These results suggest that a post-learning sleep interval strengthens MTL–neocortical communication, thereby facilitating more efficient reactivation of associative memory traces. Sleep has been proposed to promote repeated reactivation of newly encoded information within the MTL and its gradual redistribution to long-term neocortical areas ^25,26,39,40^. Accordingly, our results revealed sleep-dependent increases in neocortical synchronization, particularly between the right inferior occipital gyrus, right inferior temporal gyrus, and bilateral IFG, possibly reflecting the progressive reorganization of memory traces toward long-term semantic storage sites. Supporting this interpretation, using the same experimental material, our group previously reported sleep-dependent increases in bilateral IFG evoked magnetic responses within the same time window (450–650 ms) in children, interpreted as evidence for rapid consolidation within episodic–semantic networks ^42^. Still, sleep-dependent increases in theta-band synchronization remained anchored in MTL structures such as the left parahippocampal, entorhinal, and perirhinal cortices, suggesting that sleep strengthens MTL–neocortical interactions rather than promoting MTL disengagement. This pattern is consistent with the Multiple Traces Theory, which posits a sustained role of MTL structures in the recall of detail-rich memories across development ^62,63^. Interestingly, the left postcentral gyrus also emerged as a central connectivity hub in our results, interacting with both MTL structures and other neocortical regions known to be implicated in semanticization processes, such as the right temporal pole ^58^. These results align with sensorimotor-based theories of semantic memory which posit that semantic information about an object is partly grounded in neural systems involved in perception and action ^64^. This further suggests that sleep may facilitate the integration of newly learned function–object associations into sensorimotor–semantic networks. In this perspective, the sleep-dependent MTL–neocortical network observed in our results, also aligns with recent fMRI or EEG studies suggesting that offline consolidation episodes, involving sleep, possibly influence the transformation of memory traces into higher-order semantic (i.e., gist-based) representations and/or indicate post-sleep changes in recall strategies ^20,65^. Yet, these studies did not investigate the specific impact of sleep on these processes as, contrary to our study, they did not control that these processes were not occurring after a similar interval of offline wakefulness.

Moreover, to further investigate how sleep-specific oscillatory activity supports memory recall, we examined the neural mechanisms underlying delayed recall within the Sleep group. Transient theta-band phase synchronization in the 450–650 ms post-cue window involved a predominantly left-lateralized network, primarily involving MTL– neocortical connections, corroborating our previous findings that sleep (relative to wakefulness) enhances MTL–neocortical communication. Critically, the mean network synchronization strength in this post-sleep network, as well as memory performance gains, were positively associated with slow-oscillation–spindle PAC during the nap, whereas global sleep measures were not. This finding builds on prior evidence that slow-oscillation–spindle coupling mediates successful memory consolidation ^34–37^ and provides evidence that this mechanism reinforces MTL–neocortical communication during delayed recall, potentially facilitating more efficient reactivation of associative memory traces. Together, these results extend established systems consolidation models to childhood, demonstrating that sleep-dependent plasticity mechanisms, such as slow-oscillation–spindle PAC, are already operative in the developing brain and may even be particularly effective, as indicated by their observable effects after a single nap.

Our study has several limitations. First, group differences in theta-band phase synchronization were observed in the absence of corresponding differences in memory performance gains. However, this is consistent with previous neuroimaging studies that reported sleep-related neural reorganization without concomitant behavioral change ^42,66,67^. This dissociation may reflect ceiling effects induced by the learning criterion ^68^, or the possibility that longer retention intervals and additional nights of sleep are required for sleep-dependent behavioral benefits to emerge reliably ^69,70^. Second, although theta-band synchronization around 500 ms is consistent with cortical reinstatement of memory traces, we could not directly demonstrate reactivation of encoding-related neuronal ensembles, as MEG data were not acquired during encoding phase. Future studies combining encoding– retrieval recordings or multivariate pattern analyses could provide more direct evidence for reactivation mechanisms ^71,72^. Finally, developmental comparisons between children and adults will be necessary to determine whether these effects genuinely reflect heightened plasticity during childhood and the cognitive advantages of this sensitive period.

In summary, this study reveals a temporally ordered sequence of theta-band brain synchronization dynamics underlying memory recall in children, progressing from an early visual pathway–hippocampal network to a later, large-scale MTL–neocortical network, consistent with memory reinstatement processes. We further demonstrate that even a brief daytime nap can selectively enhance theta-band synchronization, particularly between MTL and neocortical regions during delayed recall, likely via sleep-specific plasticity mechanisms, thereby supporting a more efficient reinstatement of associative memory traces in the developing brain.

## 4. Methods

### 4.1. Participants

Forty-one 7–12 years old typically developing children were randomly assigned to a Sleep (N=23) or a Wake (N=18) condition. A written informed consent was given by all participants and their parents to participate in this MEG study. The study protocol was approved by the Ethics Committee of the Hôpital Universitaire de Bruxelles (HUB) - Hôpital Erasme (Brussels, Belgium; Reference: P2018/335). All children were native French speakers with self-reported normal or corrected vision. They had no prior history of neurological or psychiatric issues neither of any learning, cognitive or language disabilities. Based on the Sleep Disturbances Scale for Children” (SDSC) ^73^, all children had normal sleep quality and sleep habits over the 6 months preceding the experiment (SDSC total score < 67). We additionally ensured that they had maintained a regular sleep schedule for the 3 nights preceding the experiment using the “St Mary’s Hospital Sleep Questionnaire” ^74^ (see Supplementary Information, SI8). Overall, 10 children were excluded from the analyses either due to excessive movements during MEG recordings (N=5), poor memory recall performance (more than 2.5 standard deviations; N=2) or, in the Sleep group, lack of NREM stage 3 sleep in the post-learning nap (N=3). The final sample was consequently composed of 31 children (15 girls and 16 boys, mean age ± SD: 9.90 ± 1.27 years; range, 7 to 12 years), including 15 children in the Sleep group (8 girls and 8 boys; mean age ± SD: 9.80 ± 1.15 years) and 16 children in the Wake group (7 girls and 8 boys; mean age ± SD: 10.00 ± 1.41 years). The Sleep and Wake groups were matched for age (independent-samples *t*-test; *p* = 0.67) and sex (χ^2^ test; *p* = 0.59).

### 4.2. Material and design

#### 4.2.1. Material

A detailed description of the material and behavioral learning task can be found in ^75^. Briefly, one hundred 2D colored drawings of unknown non-objects (Figure 4A) were matched using Photoshop− for contrast, brightness, size (6×6 cm on screen), and color. Four lists of 50 to-be-learned non-objects were created, and one list was assigned to each child during the learning phase, in a counterbalanced order. For each list, the remaining 50 non-objects served as control non-objects. Each to-be-learned non-object was randomly paired with a magical (imaginary) function that participants had to learn (“*With this object you can*”: e.g., “paint the sky in colors”, “open any door”, “stop the rain”). Each function was presented to the child in French, using sentences ranging from 3 to 7 words in length.

**Figure 4.**
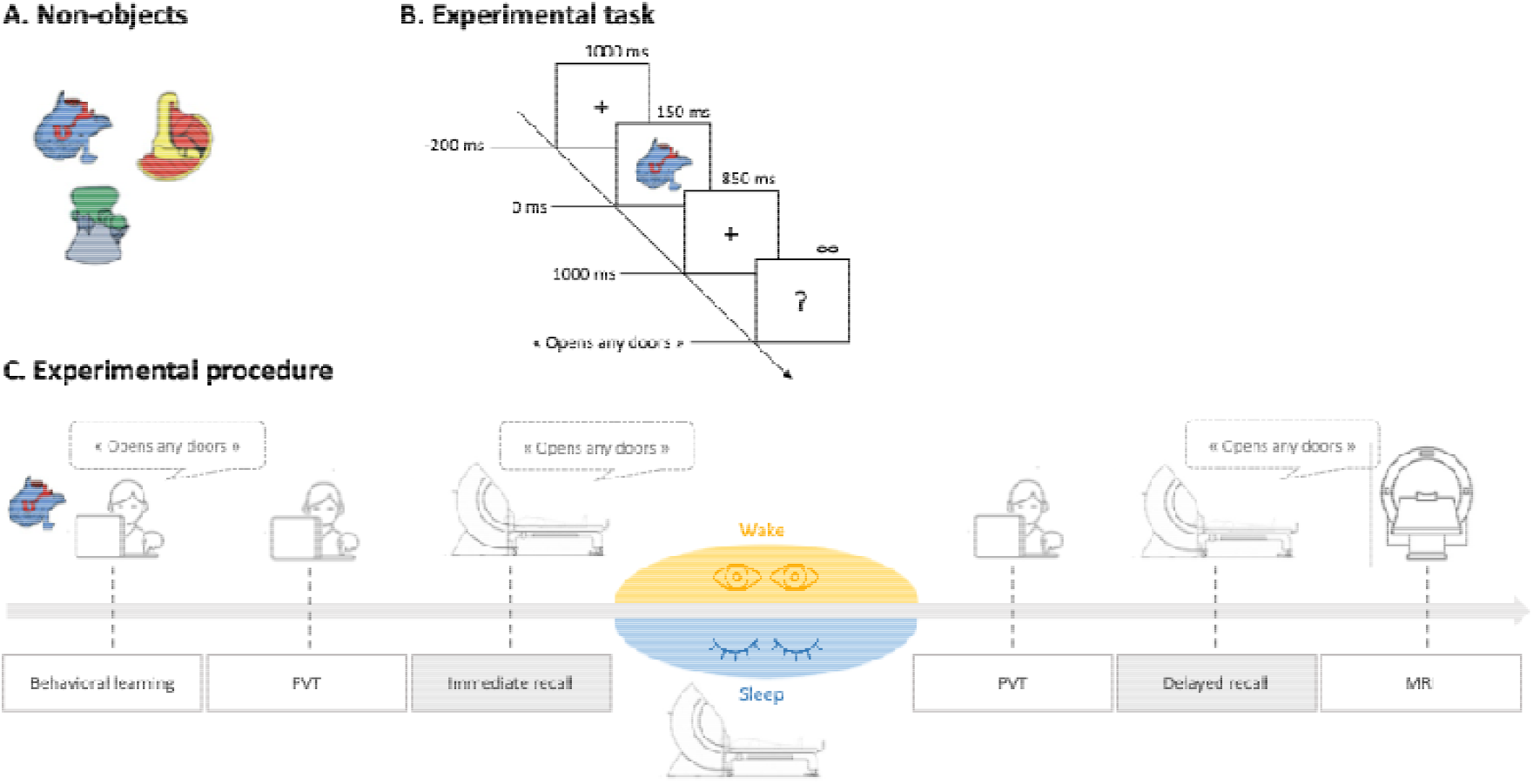
Experimental task and procedure. **(A)** Sample illustrations of the 50 non-objects used. **(B)** Experimental memory task: At each trial, children were asked to provide the function associated with the non-object presented on the screen. Responses had to be given after the appearance of the question mark (1 s after cue onset). **(C)** Experimental procedure: After the behavioral learning session, children undergo two MEG sessions (immediate recall and delayed recall), preceded by a PVT and separated by either a nap (Sleep group) or a wake (Wake group) period. A final structural high-resolution brain 3D T1-weighted magnetic resonance image is acquired.

#### 4.2.2. Experimental task and procedure

This study occurred in the context of a one-day protocol (Figure 4C) starting in the morning with the behavioral learning session requiring each child to learn, outside the MEG scanner, the associations between the 50 to-be-learned non-objects and their magical functions. Participants had to reach a specific 60% learning criterion (30 of 50 items mastered), ensuring sufficient learned stimuli for MEG data analysis while keeping children motivated during the learning session and allowing for sufficient interindividual variability in scores. If the participant did not reach 60% correct responses on the first attempt, the learning session was repeated with presentation of the unlearned non-objects only. This behavioral learning session was followed by two MEG recall sessions: an immediate recall session and a delayed recall session, during which children were asked to retrieve the functions associated with the 50 non-objects. The two sessions were strictly similar. Stimuli appeared on a projection screen located 1 m in front of the child who was lying in the MEG scanner to prevent excessive motions. Children were asked to retrieve the functions associated with the 50 to-be-learned non-objects and to correctly skip the 50 control non-objects. Each set of 50 non-objects was presented twice in a random order. During each trial, a non-object cue was presented for 150 ms, followed by an 850 ms blank screen with a fixation cross and then a question mark prompting the child to verbally provide aloud the object’s function or say “skip” if unknown. The question mark remained on the screen until a response was given, and the next trial began after a 1000 ms inter-stimulus interval corresponding to a blank screen with a fixation cross (Figure 4B). The delayed definition response procedure we chose (as opposed to an immediate verbal response that would have allowed the measurement of reaction times) equalizes as much as possible the process of word retrieval between children and reduces mouth movement artifacts in MEG recording during the post-cue period of interest.

The two MEG recall sessions were separated by either a 90-minute interval of quiet wakefulness (Wake Group) during which children were asked to calmly engage in familiar activities outside the MEG scanner, or a 90-minute nap in the MEG aimed at obtaining at least one full sleep cycle (Sleep Group), both occurring after lunchtime under controlled conditions.

To ensure that experimental effects could not be attributed to variations in vigilance between groups (Sleep vs. Wake) and sessions (immediate vs. delayed recall), vigilance performance was also measured before each MEG recall session using the PVT (5-minute version) during which participants were instructed to press a response button as quickly as possible whenever a digital counter stimulus appeared ^76^. The protocol ended with the acquisition of a structural high-resolution 3D T1-weighted brain magnetic resonance image (MRI) which allowed MEG source reconstruction through the coregistration of functional MEG data with individual structural images for each participant (see below).

### 4.3. MEG and MRI data acquisition

Participants’ MEG data (signals band-pass filtered at 0.1–330 Hz and sampled at 1 kHz) were recorded inside a magnetically shielded room (Maxshield, MEGIN, Helsinki, Finland; see ^77^ for details) using a 306-channel whole-scalp neuromagnetometer (MEG) system (Triux, MEGIN, Helsinki, Finland) installed at the H.U.B–Hôpital Erasme. Throughout the run, children head position in the MEG helmet was continuously monitored with four head-tracking coils ^78^. Three landmark positions (left and right tragi and nasion) and at least 400 additional head-surface points (on scalp, nose, and face) were digitized using an electromagnetic tracker system (Fastrak, Polhemus, Colchester, VT, USA). MEG-compatible bipolar electrodes were used to monitor ocular, cardiac, and mouth muscle artefacts, positioned vertically around the eyes (electrooculography, EOG), on the back (electrocardiography, ECG), and vertically on the chin (electromyography, EMG). Sleep was recorded simultaneously with MEG and using 5 MEG-compatible EEG electrodes (F3, F4, C3, C4 and CZ) referenced to the opposite mastoid.

After the MEG sessions, high-resolution anatomical 3D T1-weighted MRI scans were acquired (MRI, 1.5T, Intera, Philips, Best, The Netherlands) in all participants except in 7 children for whom we used a linear deformation of the structural MRI of other age-matched children to best match head-surface points, using the CPD toolbox ^79^ embedded in FieldTrip (Donders Institute for Brain Cognition and Behaviour, Nijmegen, The Netherlands, RRID:SCR_004849) ^80^.

### 4.4. Behavioral analyses

Behavioral data analyses were performed using Jamovi (Version 2.3) ^81^. Memory recall performance was calculated for each participant as the percentage of correctly recalled associations (i.e., to-be-learned non-object–function) during the immediate and delayed recall sessions. Outliers were defined as recall performance values exceeding 2.5 standard deviations from the session-specific mean and were removed from the analyses. To assess time-dependent changes in memory performance and the specific effect of sleep (vs. wakefulness), we conducted a mixed-design ANOVA with session (immediate vs. delayed recall) as a within-subject factor and group (Sleep vs. Wake) as a between-subject factor. For the sake of completeness, we additionally conducted a complementary Generalized Linear Mixed Model (GLMM) analysis on recall performance, examining the effects of session (immediate vs. delayed recall), group (Sleep vs. Wake), and the interaction between group and session using RStudio (Version 2023.12.0) ^82^ with R (Version 4.3.2) ^83^. This approach allowed us to analyze participants’ memory recall performance while accounting for both inter-trial (by-item random intercepts) and interindividual variability (random intercepts and slopes for session by participant) (see more details regarding this analysis methods and results in Supplementary Information, SI9). Memory gain was also computed as the percentage change in performance between the immediate and delayed recall sessions using the following formula: ((delayed recall –immediate recall) / immediate recall) x100. Positive values indicate improved recall performance over time, whereas negative values reflect performance decline, with zero indicating no change in performance. Finally, vigilance was assessed at each recall session using mean reciprocal reaction times (RRTs=1/Reaction Time(s)) ^48^ from the PVT and analyzed using a mixed-design ANOVA with one within-subject factor session (immediate vs. delayed recall) and one between-subject factor group (Sleep vs. Wake).

### 4.5. MEG analyses

#### 4.5.1. MEG data preprocessing and source reconstruction

MEG data were filtered using a temporal extension of signal space separation (tSSS; correlation coefficient, 0.98; window length, 10s) ^78^ to remove external environmental noise and correct for head movements (Maxfilter TM, Elekta Oy, Helsinki, Finland; version 2.2 with default parameters) and band-pass filtered offline at 1–40 Hz. An independent component analysis (FastICA algorithm with dimension reduction to 30 and hyperbolic tangent non-linearity contrast) ^84^ was additionally performed on MEG signals to eliminate remaining ocular and cardiac artifacts. Components related to artifacts were visually inspected and removed from the full rank data for each MEG session (mean number of components ± SD for immediate and delayed recall sessions: 3.6 ± 0.8, range: 2–5 and MEG sleep: 1.8 ± 0.7; range 1–4). Individual brain MRI was anatomically segmented using FreeSurfer software (Martinos Center for Biomedical Imaging, Massachussetts, USA) ^85^ and MEG functional data and MRI structural data were manually coregistered using the digitized fiducials points and refined using the additional points on the head surface (Mrilab, MEGIN Croton Healthcare, Helsinki, Finland). A volumetric source grid (5 mm) was then established within the Montreal Neurological Institute (MNI) template MRI and deformed onto each subject’s MRI through non-linear deformation in Statistical Parametric Mapping Software (SPM12, Wellcome Trust Centre for Neuroimaging, London, UK) ^86^. The MEG forward model was subsequently computed using the Boundary Element Method, implemented in the MNE-C suite (Martinos Center for Biomedical Imaging, Massachusetts, USA) ^87^.

Neuronal source activity was then reconstructed using a Minimum Norm Estimation model (MNE) ^88^ within the theta (i.e., 4–8 Hz) frequency band due to the strong theoretical *a priori* related to the central role of theta oscillations in memory encoding and recall processes ^17–19,89^.

A 10-minute MEG empty room recording was also filtered in the specified frequency band and preprocessed with SSS to estimate the noise covariance matrix. The MNE regularization parameter was determined based on the consistency condition outlined in a previous study ^90^. Resulting three-dimensional dipole time series were then analyzed by projecting them onto their direction of maximum variance and obtaining their analytic signal through Hilbert transformation, following the methodology previously described ^91,92^.

#### 4.5.2. Theta-band phase synchronization analyses related to immediate and delayed memory recall

##### Theta-band phase synchronization analyses

Instantaneous theta phase of source signals associated with correctly recalled associations (i.e., non-object–function) was extracted using Hilbert transform (Figure 5A), epoched from 200 ms prior to 1100 ms after cue onset (i.e., non-object presentation), and finally, downsampled every 20 ms. To unfold the dynamics of theta-band phase synchronization associated with the recall of non-object–function associations, we computed the cross-trial phase locking value (PLV; Figure 5B). Ranging between 0 and 1, the PLV reflects phase synchrony between two sources, which is understood to be a neurophysiological mechanism mediating communication among brain regions ^93^. This measure of synchronization was computed between all pairwise combinations of 75 nodes ^94^ (see Supplementary Information, SI10 for MNI coordinates), based on a customized parcellation of the human brain adapted from the Automated Anatomical Labeling (AAL) atlas ^95^, and including specific declarative memory nodes described in previous studies ^40,42,53,96^. Spatial leakage was corrected between each node pair using the geometric correction scheme ^90^. The resulting 75-by-75 matrices of theta-band PLV were calculated for each subject and time step (from - 200 to 1100 ms, step every 20 ms; C 5). To control for inter-subject variability in resting-state theta phase synchrony, PLV time courses were z-scored over baseline time samples (-200 to 0 ms).

**Figure 5.**
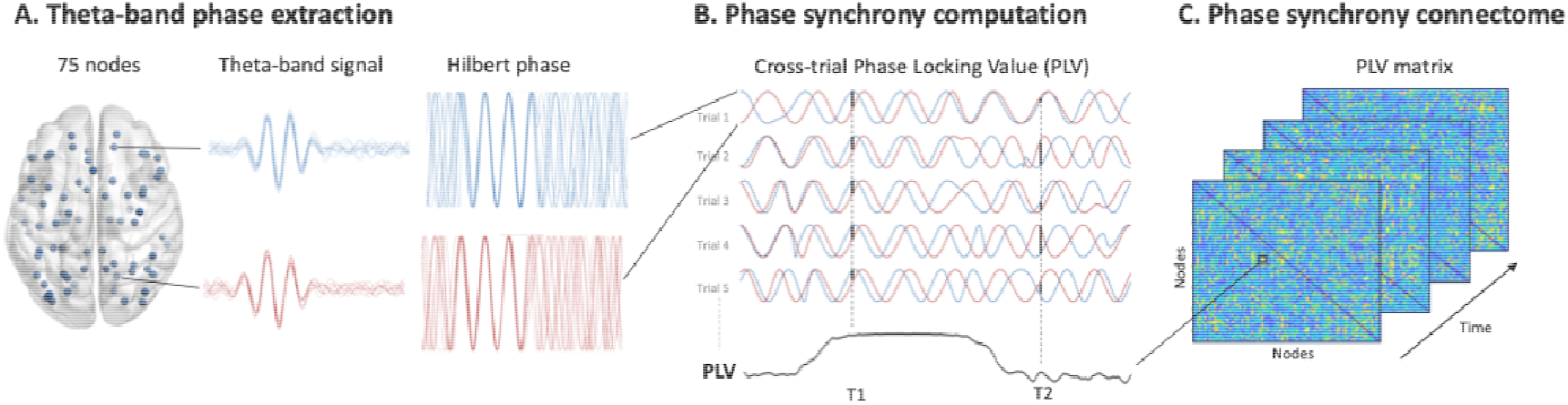
Schematic overview of the theta-band phase synchronization analyses. **(A)** Following MEG data preprocessing and source reconstruction, instantaneous theta-band signals were extracted from 75 brain regions (i.e., nodes), and their phase time courses were obtained using the Hilbert transform. The illustrated signals correspond to simulated, epoched data containing time-locked transient events of theta-band phase synchronization, resulting in a visible theta oscillation in band-pass–filtered signals and to synchronization in a window in the corresponding Hilbert phase time courses. **(B)** Theta-band phase synchrony between pairs of nodes was quantified using the cross-trial phase-locking value (PLV), which assesses the consistency of phase differences at each time point across repeated trials. As illustrated, a consistent phase lag across trials at time point T1 yields a high PLV, whereas variable phase lags across trials at time point T2 result in a low PLV. **(C)** This procedure was applied to all pairs of regions, producing time-resolved 75-by-75 theta-band PLV matrices for each participant and time point, which were subsequently used for network analyses.

##### Statistical analyses

To characterize the temporal dynamics of theta-band phase synchronization, z-scored PLV were averaged across source pairs and subjects and plotted over each time point. This provided a global, whole-brain view of interregional theta-band PLV dynamics, allowing us to identify time windows of interest for subsequent network analyses without spatial priors. To identify significantly synchronized network, PLV matrices (75-by-75 nodes) associated with selected time windows were analyzed using the NBS software ^49,50^. Statistical contrasts were computed either between conditions (selected time window vs. baseline) or between groups (Sleep vs. Wake). NBS is a non-parametric statistical tool which performs multiple univariate tests on all analyzed edges (in this case each element in the PLV matrix; see similar approaches ^97–99^), while controlling for multiple comparisons across the connectome ^49,50^.

The primary component-forming threshold was adapted to the distribution of the data so as to retain 1% of total possible network connections (≈28 edges out of 2,775), following previous work using a similar approach ^100^. Statistical significance of the resulting network components was assessed using 10,000 permutations, with a corrected significance threshold set at p^corr^□<□0.05. For each significant network identified through NBS, node-level contributions were quantified using a node strength measure, defined as the sum of all weighted edges connected to a given node ^101^. The resulting brain networks and weighted nodes were visualized using the BrainNet Viewer Connectivity Toolbox ^102^. The mean network synchronization strength was also calculated for each participant as the averaged node strengths in the significant network. Supplementary one-tailed Pearson’s correlations were computed to investigate associations between mean network synchronization strength and memory recall performance.

#### 4.5.3. Associations with slow-oscillation–spindle phase amplitude coupling during sleep

Sleep EEG data were first preprocessed and analyzed by a sleep scoring expert (PS) using the Fercio’s software (FercioEEGPlus, Ferenc Gombos, 2008–2018). Sleep stages were manually scored according to standardized criteria (AASM) ^103^ on 20 s epochs. Total sleep time, as well as sleep stages duration and percentage were extracted (see Supplementary Information, SI7). Slow-oscillation–spindle PAC was then computed from sleep MEG signals restricted to NREM stages 2 and 3 based on sleep scoring of simultaneous EEG. NREM stages 2 and 3 were selected to optimize slow-oscillation–spindle coupling detection as stage 2 is characterized by frequent sleep spindles, whereas stage 3 exhibits prominent slow-oscillation activity. Spindle activity was quantified as amplitude variation within the sigma frequency range (11–15 Hz) ^104,105^. Individual spindle events were not analyzed in this study. PAC was computed within each of the 102 gradiometer pairs using the modulation index (MI) ^106^. The MI quantifies phase-amplitude coupling between a phase-modulating frequency range and an amplitude-modulated frequency range. The frequency for phase was varied from 1 to 8 Hz by steps of 1 Hz, with a 1 Hz bandwidth, and the frequency for amplitude was varied from 10 to 40 Hz by steps of 2 Hz, with a bandwidth set to 30% of the center frequency. MEG gradiometer signals were filtered within these frequency bands, and gradiometer pairs were combined using the first principal component. Thereafter, the Hilbert transform was applied to extract the instantaneous phase time series of the signals for phase, and the amplitude envelope of the signals for amplitude. For each pair of frequency for phase and frequency for amplitude, the phase values were divided into 18 bins, and the mean amplitude was computed for each phase bin. The resulting amplitude distribution was normalized by dividing each mean amplitude by the sum across all bins, yielding a probability distribution. MI was then derived using the Kullback–Leibler divergence to quantify the deviation from a uniform distribution ^106^. MI values close to 0 indicate no coupling (a nearly uniform amplitude distribution), while higher values suggest stronger phase-dependent modulation of higher frequency amplitude by slower oscillations. Further analyses were conducted on the slow-oscillation–sigma PAC value, which was the MI at the frequency for phase centered on 1 Hz averaged across the frequency for amplitude of 12 and 14 Hz (which approximate the 11–15 Hz sigma frequency band). For each participant, a relative slow-oscillation–sigma PAC value was computed as the maximum slow-oscillation– sigma MI across gradiometer pairs divided by its median. This provided a participant-specific PAC measure that is minimally affected by the amount of data used to derive it. Finally, Pearson correlation analyses were conducted to investigate the associations between slow-oscillation–spindle PAC and the quality of delayed memory recall processes quantified as the behavioral gain in performance and the mean network synchronization strength in the sleep group.

## Data availability

Code and processed data underlying the results will be made available upon request to the corresponding author or will be made available in a repository if requested.

## Acknowledgements and funding

SG was supported by the Fonds pour la formation à la recherche dans l’industrie et l’agriculture (FRIA, Fonds de la Recherche Scientifique (FRS-FNRS), Brussels, Belgium) and by a research Seed Money from the Université libre de Bruxelles (Brussels, Belgium). XDT is Clinical Researcher at the Fonds de la Recherche Scientifique (FRS-FNRS, Brussels, Belgium). The MEG project at the Hôpital Universitaire de Bruxelles and Université libre de Bruxelles is financially supported by the Fonds Erasme (Convention « Les Voies du Savoir », Brussels, Belgium) and by research grants from the FRS-FNRS (FRS-FWO Excellence Of Science (EOS) MEMODYN: 30446199 ; CREDIT DE RECHERCHE: n° 29149840). The authors would also like to warmly thank all the children and their parents for their participation.

## Authors contributions

S.G.: behavioral, MEG, EEG and MRI data analyses, interpretation of the results, writing - original draft. C.R.: MEG and MRI data analyses, revising the manuscript. A.P.: data acquisition, revising the manuscript. P.P.: conceptualization, interpretation of the results, revising the manuscript, funding. P.S.: EEG data analyses, interpretation of the results, revising the manuscript. M.B.: software, MEG data analyses, interpretation of the results, revising the manuscript. V.W.: software, MEG, and MRI data analyses, interpretation of the results, writing - original draft. X.D.T.: conceptualization, interpretation of the results, revising the manuscript, funding. G.D.: behavioral data analyses, interpretation of the results, writing - original draft, funding. C.U.: conceptualization, interpretation of the results, writing - original draft, funding.

