## Supplementary Information 1 for "Theta-band brain synchronization supports the immediate and post-sleep dynamics of memory recall in children"

### Supplementary Information 1

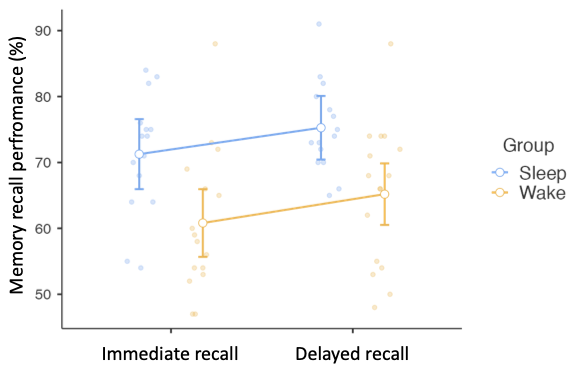

***Figure S1.*** ***Memory recall performance across sessions and groups.*** *Memory recall performance in the Sleep and Wake groups during immediate and delayed recall sessions. Data were analyzed using a mixed ANOVA with session (immediate vs. delayed) as a within-subject factor and group (Sleep vs. Wake) as a between-subject factor. Error bars represent 95% confidence intervals.*

### Supplementary Information 2

| X | Y | Z | Node strength | Label |
| --- | --- | --- | --- | --- |
| 42 | -56 | -10 | 17,01 | Temporal_Inf_R |
| 24 | -81 | 31 | 15,07 | Occipital_Sup_R |
| 16 | -73 | 9 | 12,50 | Calcarine_R |
| -20 | -63 | 17 | 9,06 | Calcarine_L |
| 16 | -67 | -4 | 8,56 | Lingual_R |
| -32 | -81 | 16 | 7,77 | Occipital_Mid_L |
| 10 | -56 | 44 | 5,71 | Precuneus_R |
| -15 | -68 | -5 | 5,54 | Lingual_L |
| 30 | -58 | 26 | 4,84 | Occipital-parietal_R |
| -36 | -78 | -8 | 4,63 | Occipital_Inf_L |
| 40 | -74 | 34 | 3,05 | Parietal_Inf_R |
| 31 | -53 | -2 | 2,86 | Fusiform_R |
| -5 | -43 | 25 | 2,77 | Cingulum_Post_L |
| 18 | -62 | 15 | 2,28 | Calcarine_R |
| -11 | 11 | 9 | 2,23 | Caudate_L |
| 38 | -82 | -8 | 2,14 | Occipital_Inf_R |
| 7 | -42 | 22 | 2,07 | Cingulum_Post_R |
| -29 | -21 | -14 | 2,01 | HPC_L |
| -23 | -60 | 59 | 1,91 | Parietal_Sup_L |
| -50 | -28 | -23 | 1,74 | Temporal_Inf_L |
| 16 | -52 | 8 | 1,55 | Retrosplenial_R |
| -23 | -1 | -17 | 1,50 | Amygdala_L |

***Table S1.*** ***Nodes of the significant early theta synchronization network (150–350 ms).*** *List of brain regions showing significant theta-band phase synchronization relative to baseline (−200 to 0 ms) during immediate recall in the early post-cue window (150–350 ms). Node strength and MNI coordinates (mm) are reported for each region.*

### Supplementary Information 3

| X | Y | Z | Node strength | Label |
| --- | --- | --- | --- | --- |
| 30 | -58 | 26 | 5,62 | Occipito-parietal_R |
| -15 | -68 | -5 | 4,69 | Lingual_L |
| 54 | -31 | -22 | 3,85 | Temporal_Inf_R |
| 44 | 15 | -32 | 3,72 | Temporal_Pole_Mid_R |
| -23 | -1 | -17 | 3,33 | Amygdala_L |
| 52 | 20 | 8 | 2,82 | IFG_R |
| 38 | -82 | -8 | 2,74 | Occipital_Inf_R |
| 42 | -56 | -10 | 2,42 | Temporal_Inf_R |
| 28 | -10 | -37 | 2,39 | Entorhinal_R |
| -5 | -43 | 25 | 2,22 | Cingulum_Post_L |
| -35 | 7 | 3 | 2,13 | Insula_L |
| -23 | -60 | 59 | 2,10 | Parietal_Sup_L |
| 38 | 33 | 34 | 1,99 | Frontal_Mid_R |
| -24 | -40 | -12 | 1,81 | PHPC_L |
| 57 | -37 | -1 | 1,46 | Temporal_Mid_R |
| 0 | 20 | 64 | 1,35 | Frontal_Medial_Post_R |
| 21 | -19 | -23 | 1,21 | PHPC_R |
| 38 | 2 | -40 | 1,17 | Temporal_Inf_R |
| 27 | 1 | -18 | 1,05 | Amygdala_R |
| -24 | -13 | -37 | 1,03 | Entorhinal_L |
| 31 | -53 | -2 | 0,69 | Fusiform_R |

***Table S2. Nodes of the significant late theta synchronization network (550–750 ms).*** *List of brain regions showing significant theta-band phase synchronization relative to baseline (−200 to 0 ms) during immediate recall in the late post-cue window (550–750 ms). Node strength and MNI coordinates (mm) are reported for each region.*

### Supplementary Information 4

| X | Y | Z | Node strength | Label |
| --- | --- | --- | --- | --- |
| -38 | -68 | -16 | 7,96 | Fusiform_L |
| -42 | -23 | 49 | 6,48 | Postcentral_L |
| -24 | -20 | -28 | 5,59 | Perirhinal_L |
| 30 | -58 | 26 | 4,69 | Occipito-parietal_R |
| 31 | -53 | -2 | 4,49 | Fusiform_R |
| -29 | -21 | -14 | 4,48 | HPC_L |
| 10 | -56 | 44 | 4,28 | Precuneus_R |
| -50 | -28 | -23 | 3,68 | Temporal_Inf_L |
| 16 | -52 | 8 | 3,56 | Retrosplenial_R |
| -21 | -19 | -23 | 3,01 | PHPC_L |
| -42 | -42 | 16 | 2,72 | Temporal_Sup_L |
| -31 | -40 | -20 | 2,08 | Fusiform_L |
| -11 | 11 | 9 | 1,80 | Caudate_L |
| -23 | -1 | -17 | 1,77 | Amygdala_L |
| -35 | 7 | 3 | 1,71 | Insula_L |
| -24 | -40 | -12 | 1,71 | PHPC_L |
| 36 | -12 | 70 | 1,68 | Frontal_Sup_R |
| -24 | -13 | -37 | 1,44 | Entorhinal_L |
| 32 | -20 | -13 | 1,41 | HPC_R |
| -39 | -6 | 51 | 1,27 | Precentral_L |
| 16 | -67 | -4 | 1,26 | Lingual_R |
| -48 | 13 | 19 | 1,10 | Frontal_Inf_Oper_L |
| -44 | -61 | 36 | 0,93 | Angular_L |

***Table S3. Nodes of the significant theta synchronization network showing a group effect (Sleep > Wake).*** *List of brain regions showing significant theta-band phase synchronization differences between groups (Sleep > Wake) during delayed recall in the post-cue window (450–650 ms). Node strength and MNI coordinates (mm) are reported for each region.*

### Supplementary Information 5

| X | Y | Z | Node strength | Label |
| --- | --- | --- | --- | --- |
| -38 | -68 | -16 | 7,96 | Fusiform_L |
| -42 | -23 | 49 | 6,48 | Postcentral_L |
| -24 | -20 | -28 | 5,59 | Perirhinal_L |
| 30 | -58 | 26 | 4,69 | Occipito-parietal_R |
| 31 | -53 | -2 | 4,49 | Fusiform_R |
| -29 | -21 | -14 | 4,48 | HPC_L |
| 10 | -56 | 44 | 4,28 | Precuneus_R |
| -50 | -28 | -23 | 3,68 | Temporal_Inf_L |
| 16 | -52 | 8 | 3,56 | Retrosplenial_R |
| -21 | -19 | -23 | 3,01 | PHPC_L |
| -42 | -42 | 16 | 2,72 | Temporal_Sup_L |
| -31 | -40 | -20 | 2,08 | Fusiform_L |
| -11 | 11 | 9 | 1,80 | Caudate_L |
| -23 | -1 | -17 | 1,77 | Amygdala_L |
| -35 | 7 | 3 | 1,71 | Insula_L |
| -24 | -40 | -12 | 1,71 | PHPC_L |
| 36 | -12 | 70 | 1,68 | Frontal_Sup_R |
| -24 | -13 | -37 | 1,44 | Entorhinal_L |
| 32 | -20 | -13 | 1,41 | HPC_R |
| -39 | -6 | 51 | 1,27 | Precentral_L |
| 16 | -67 | -4 | 1,26 | Lingual_R |
| -48 | 13 | 19 | 1,10 | Frontal_Inf_Oper_L |
| -44 | -61 | 36 | 0,93 | Angular_L |

***Table S4. Nodes of the significant theta synchronization network in the Sleep group (450–650 ms).*** *List of brain regions showing significant theta-band phase synchronization relative to baseline (−200 to 0 ms) during delayed recall in the post-cue window (450–650 ms) for the Sleep group. Node strength and MNI coordinates (mm) are reported for each region.*

### Supplementary Information 6

We further qualitatively characterized whether post-sleep increase in theta-band synchronization in the 450–650 ms post-cue window at delayed recall involved new and/or pre-existing connections (edges) as compared with the immediate recall (450–650 ms) (Figure S2A). This descriptive analysis showed that while some brain regions (nodes) overlapped between the immediate and delayed recall networks in the sleep group (i.e., orange nodes), none of the connections (edges) identified in the post-sleep delayed recall (i.e., blue edges) network were present in the immediate recall network (i.e., red edges, see Figure S2B). Furthermore, several nodes involved in the post-sleep delayed recall network including the bilateral hippocampal areas and left MTL regions (e.g., the parahippocampal, entorhinal and perirhinal cortices) as well as the left inferior and superior temporal gyrus, were absent from the immediate recall network at the same time window (450–650 ms post-cue window).

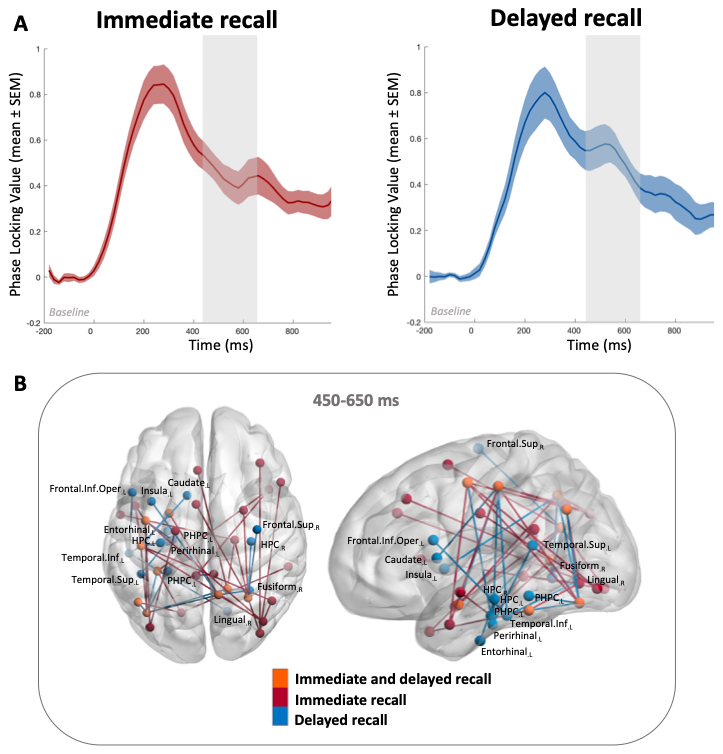

***Figure S2****.* ***Theta-band synchronization dynamics during memory recall in the sleep group.*** ***(A)*** *Theta-band phase synchronization computed as cross-trial PLV (mean ± SEM) in the Sleep group during immediate (left) and delayed (right) recall.* ***(B)*** *Brain regions forming significant synchronized networks in the* *450–650 ms post-cue window compared to baseline (-200–0 ms) during immediate recall (red nodes), delayed recall (blue nodes) or both sessions (orange nodes). Regions specific to the delayed post-sleep recall network are labelled on the brain plot.*

### Supplementary Information 7

During the post-learning nap occurring in-between the immediate and the delayed recall, children from the Sleep group slept on average 67 ± 21 minutes, including 29 ± 10 minutes of stage 2 and 23 ± 9 minutes of stage 3 (i.e., slow wave sleep) NREM sleep for a total recording time of 94 ± 14 minutes. Polysomnographic values are reported in Table S5. Neither total sleep duration nor the duration and percentage of individual sleep stages was significantly correlated with mean network synchronization strength of the post-sleep delayed recall network or memory performance gains (all *ps* > 0.5).

|  | Duration [min] | Relative duration [% of total record time] |
| --- | --- | --- |
| Total recording time | 94 ± 14 |  |
| Total sleep time | 67 ± 21 | 71 ± 15 |
| Total wake time | 27 ± 13 | 29 ± 15 |
| Stage 1 sleep | 6 ± 4 | 10 ± 5 |
| Stage 2 sleep | 29 ± 10 | 44 ± 10 |
| Stage 3 sleep | 23 ± 9 | 35 ± 11 |
| REM sleep | 9 ± 8 | 11 ± 10 |

***Table S5.* *Post-learning nap parameters.*** *Mean ± SD values are presented for the total recording time, total sleep time and time spent awake. The second part of the table report time spent in each sleep stage (i.e., NREM stage 1, stage 2 and stage 3, and REM sleep), reported in minutes and as a percentage of total sleep time.*

### Supplementary Information 8

According to parental reports using the Sleep Disturbances Scale for Children” (SDSC) ^1^ and the “St Mary's Hospital Sleep Questionnaire” ^2^, children had normal sleep quality (mean SDSC score ± SD: 36.0 ± 5.5) and maintained a regular sleep schedule in the days preceding the experiment. Indeed, average sleep duration over the past six months evaluated with the SDSC (mean ± SD: 9.7 ± 0.6 h) did not differ from the reported sleep duration for the three nights before the experiment (night −3: 9.9 ± 0.8 h; night −2: 9.9 ± 0.9 h; night −1: 9.6 ± 0.7 h; p = 0.22). Importantly, sleep duration and sleep quality did not differ between the Sleep and Wake groups (SDSC score: p = 0.06; sleep hours: all ps > 0.05.

### Supplementary Information 9

For the sake of completeness, in addition to the mixed-design ANOVA conducted on memory recall performance, we conducted a complementary Generalized Linear Mixed Model (GLMM) analysis using RStudio (Version 2023.09.1, Posit Team, 2025) with R (Version 4.3.2., R Core Team, 2023). This approach allowed us to analyse participants’ memory recall performance while accounting for both inter-trial (by-item random intercepts) and interindividual variability (random intercepts and slopes for session by participant). Responses for the immediate and delayed recall were coded as “1” for correctly recalled non-object–function associations or “0” for incorrect. Accuracy (1 = correct, 0 = incorrect) was analyzed using GLMM with a logit link-function using the ‘lme4′package ^5^. The random effect’s structure was determined by identifying the maximal random effect’s structure justified by the design ^6^. Significance of the fixed effects was assessed by performing stepwise likelihood ratio tests, in which a more complex model is compared with a simpler model, as well as an examination of Akaike Information Criterion (AIC) value, which measure the goodness of fit of the model to the data while considering its complexity. The initial model (macc0) served as the baseline model, consisting of a random effect structure including the by-participant random slop for sessions and by-items random intercepts. We then compared this model to three other models: macc1, macc2, and macc3. Macc1 includes the effect of session (immediate vs. delayed recall), while macc2 additionally adds the effect of group (Sleep vs. Wake) on accuracy. Finally, macc3 includes an interaction between session and group to assess whether the effect of session varies by group.

The results of the GLMM approach are summarized in Table S6; with lower AIC values indicating a better fit of the model to the data. Notably, model 2 (i.e., macc2), which includes both session and group effects, exhibited a lower AIC compared to the initial model (macc0) consisting of a random effect structure and macc1 including the effect of session only. Additionally, the likelihood ratio Chi-squared test showed that adding the group effect (macc2) significantly improved the model fit relative to the previous model which included only the session effect (AIC = 6606.4, χ² (1) = 7.4302, p = 0.006**). However, the inclusion of the interaction between session and group in model 3 (macc3) did not significantly improve the model fit compared to macc2 (AIC = 6608.3, χ² (1) = 0.0264, p = 0.871). Consequently, Model 2 was selected as the most suitable for our analysis, striking a balance between fit to the data and model complexity. The results suggest a significant effect of group as well as an effect of session on recall accuracy but no significant interaction between session and group (Figure S3).

|  | N para | AIC | Chisq | Df | p |
| --- | --- | --- | --- | --- | --- |
| macc0 | 5 | 6621.7 |  |  |  |
| macc1 | 6 | 6611.8 | 11.885 | 1 | < 0.001 *** |
| macc2 | 7 | 6606.4 | 7.430 | 1 | 0.006 ** |
| macc3 | 8 | 6608.3 | 0.026 | 1 | 0.871 |

***Table S6. Results of the model comparison.*** *Each model is evaluated based on Akaike Information Criterion (AIC). Model 0 serves as the baseline model, including only the intercepts. Models 1, 2, and 3 incorporate additional effects of session, group, and their interaction, respectively.*

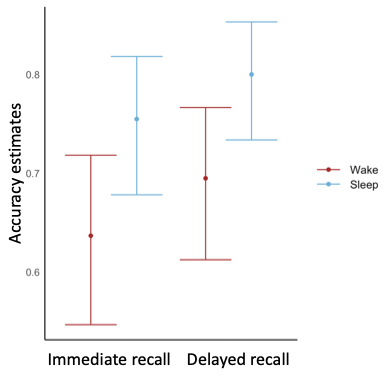

***Figure S3.*** ***Accuracy estimates from the significant model.*** *Graph generated using "ggplot2" ^7^ showing the estimated or fitted values of accuracy for each session and group from the model 2 (macc2). The error bars represent the confidence intervals around the mean of the fitted values.*

### Supplementary Information 10

| X | Y | Z | Label |
| --- | --- | --- | --- |
| 36 | -12 | 70 | Frontal_Sup_R |
| -33 | 33 | 35 | Frontal_Mid_L |
| 38 | 33 | 34 | Frontal_Mid_R |
| -48 | 13 | 19 | Frontal_Inf_Oper_L |
| 50 | 15 | 21 | Frontal_Inf_Oper_R |
| -46 | 30 | 14 | Frontal_Inf_Tri_L |
| -48 | 30 | -2 | Frontal_Inf_Tri_L |
| 50 | 30 | 14 | Frontal_Inf_Tri_R |
| 52 | 20 | 8 | Frontal_Inf_R |
| -17 | 47 | -13 | Frontal_Orb_Sup _L |
| 18 | 48 | -14 | Frontal_Orb_Sup _R |
| -36 | 31 | -12 | Frontal_Orb_Inf_L |
| 41 | 32 | -12 | Frontal_Orb_Inf _R |
| 0 | 20 | 64 | Frontal_Medial_Post_R |
| -4 | 28 | 42 | Frontal_Medial_Sup_L |
| 16 | 38 | -6 | mPFC _R |
| -16 | 38 | -6 | mPFC _L |
| -39 | -6 | 51 | Precentral_L |
| -42 | -23 | 49 | Postcentral_L |
| -5 | 5 | 61 | Supp_Motor_Area_L |
| 9 | 0 | 62 | Supp_Motor_Area_R |
| -53 | -21 | 7 | Temporal_Sup_L |
| -42 | -42 | 16 | Temporal_Sup_L |
| 57 | -37 | -1 | Temporal_Mid_R |
| -46 | 4 | -28 | Temporal_Mid_L |
| 44 | 15 | -32 | Temporal_Pole_Mid_R |
| -50 | -28 | -23 | Temporal_Inf_L |
| 54 | -31 | -22 | Temporal_Inf_R |
| 42 | -56 | -10 | Temporal_Inf_R |
| 38 | 2 | -40 | Temporal_Inf_R |
| -56 | -2 | -10 | Temporal_Medial_L |
| -4 | 35 | 14 | Cingulum_Ant_L |
| -5 | -43 | 25 | Cingulum_Post_L |
| 7 | -42 | 22 | Cingulum_Post_R |
| -29 | -21 | -14 | HPC_L |
| 32 | -20 | -13 | HPC_R |
| -24 | -40 | -12 | PHPC_L |
| -21 | -19 | -23 | PHPC_L |
| 21 | -19 | -23 | PHPC_R |
| -19 | -13 | 34 | Entorhinal_L |
| -24 | -13 | -37 | Entorhinal_L |
| 28 | -10 | -37 | Entorhinal_R |
| -24 | -20 | -28 | Perirhinal_L |
| 16 | -52 | 8 | Retrosplenial_R |
| -23 | -1 | -17 | Amygdala_L |
| 27 | 1 | -18 | Amygdala_R |
| -11 | -18 | 8 | Thalamus_L |
| 13 | -18 | 8 | Thalamus_R |
| -11 | 11 | 9 | Caudate_L |
| 15 | 12 | 9 | Caudate_R |
| -35 | 7 | 3 | Insula_L |
| 39 | 6 | 2 | Insula_R |
| -23 | -60 | 59 | Parietal_Sup_L |
| -20 | -74 | 48 | Parietal_Lobe_Sup_L |
| -43 | -46 | 47 | Parietal_Inf_L |
| -34 | -76 | 49 | Parietal_Inf_L |
| 46 | -46 | 50 | Parietal_Inf_R |
| 40 | -74 | 34 | Parietal_Inf_R |
| -44 | -61 | 36 | Angular_L |
| 46 | -60 | 39 | Angular_R |
| -7 | -56 | 48 | Precuneus_L |
| 10 | -56 | 44 | Precuneus_R |
| 30 | -58 | 26 | Occipito-parietal_R |
| 24 | -81 | 31 | Occipital_Sup_R |
| -32 | -81 | 16 | Occipital_Mid_L |
| -36 | -78 | -8 | Occipital_Inf_L |
| 38 | -82 | -8 | Occipital_Inf_R |
| -20 | -63 | 17 | Calcarine_L |
| 16 | -73 | 9 | Calcarine_R |
| 18 | -62 | 15 | Calcarine_R |
| -15 | -68 | -5 | Lingual_L |
| 16 | -67 | -4 | Lingual_R |
| -31 | -40 | -20 | Fusiform_L |
| -38 | -68 | -16 | Fusiform_L |
| 31 | -53 | -2 | Fusiform_R |

***Table S7.*** ***Nodes of the customized parcellation of the human brain****. MNI coordinates (X, Y, Z) are reported in millimeters (mm) for each region.*
